# Comprehensive Assessment of Initial Adaptation of ESBL Positive ST131 *Escherichia coli* to Carbapenem Exposure

**DOI:** 10.1101/2024.07.31.606066

**Authors:** William C. Shropshire, Xinhao Song, Jordan Bremer, Seokju Seo, Susana Rodriguez, Selvalakshmi Selvaraj Anand, An Q. Dinh, Micah M. Bhatti, Anna Konovalova, Cesar A. Arias, Awdhesh Kalia, Yousif Shamoo, Samuel A. Shelburne

**Affiliations:** Department of Infectious Diseases and Infection Control, The University of Texas MD Anderson Cancer Center, Houston, TX, USA; Department of Biosciences, Rice University, Houston, TX, USA; Department of Microbiology and Molecular Genetics, McGovern Medical School, The University of Texas Health Science Center at Houston, Houston, TX, USA; Program in Diagnostic Genetics and Genomics, The University of Texas MD Anderson Cancer Center School of Health Professions, Houston, TX, USA; Division of Infectious Diseases and Center for Infectious Diseases Research, Houston Methodist Hospital and Houston Methodist Research Institute, Houston, TX, USA; Department of Laboratory Medicine, The University of Texas MD Anderson Cancer Center, Houston, TX, USA; Department of Medicine, Weill Cornell Medical College, New York, NY, USA; Department of Genomic Medicine, The University of Texas MD Anderson Cancer Center, Houston, TX, USA

**Keywords:** Non-carbapenemase carbapenem resistance, ESBL gene amplification, experimental evolution

## Abstract

**Background:** It remains unclear how high-risk *Escherichia coli* lineages, like sequence type (ST) 131, initially adapt to carbapenem exposure in their progression to becoming carbapenem resistant.

**Methods:** Carbapenem mutation frequency was measured in multiple subclades of extended-spectrum β-lactamase (ESBL) positive ST131 clinical isolates using a fluctuation assay followed by whole genome sequencing (WGS) characterization. Genomic, transcriptomic, and porin analyses of ST131 C2/*H*30Rx isolate, MB1860, under prolonged, increasing carbapenem exposure was performed using two distinct experimental evolutionary platforms to measure fast vs. slow adaptation.

**Results:** All thirteen ESBL positive ST131 strains selected from a diverse (n=184) ST131 bacteremia cohort had detectable ertapenem (ETP) mutational frequencies with a statistically positive correlation between initial ESBL gene copy number and mutation frequency (r = 0.87, *P*-value <1e-5). WGS analysis of mutants showed initial response to ETP exposure resulted in significant increases in ESBL gene copy numbers or mutations in outer membrane porin (Omp) encoding genes in the absence of ESBL gene amplification with subclade specific associations. In both experimental evolutionary platforms, MB1860 responded to initial ETP exposure by increasing *bla*_CTX-M-15_ copy numbers via modular, insertion sequence *26* (IS*26*) mediated pseudocompound transposons (PCTns). Transposase activity driven by PCTn upregulation was a conserved expression signal in both experimental evolutionary platforms. Stable mutations in Omp encoding genes were detected only after prolonged increasing carbapenem exposure consistent with clinical observations.

**Conclusions:** ESBL gene amplification is a conserved response to initial carbapenem exposure, especially within the high-risk ST131 C2/*H*30Rx subclade. Targeting such amplification could assist with mitigating carbapenem resistance development.

## INTRODUCTION

The increased worldwide prevalence of extended-spectrum β-lactamase (ESBL) producing *Escherichia coli* (ESBL-*Ec*) are a significant contributor to *E. coli* infections leading the global cause of bacterial antimicrobial resistance (AMR) associated mortality [1]. A recent population-based United States study indicated that from 2012 to 2017 there was a 53.3% increase in ESBL producing Enterobacterales infections with 87% of that increase due to ESBL-*Ec* infections [2]. Although carbapenems are a recommended treatment option for complicated ESBL-*Ec* infections [3], ESBL-*Ec* can develop carbapenem resistance in the absence of a carbapenemase within both clinical and laboratory settings [4–7]. Non-carbapenemase producing carbapenem resistant *Escherichia coli* (non-CP-CR*Ec*), typically have increased production of ESBL enzymes concurrent with changes in outer membrane permeability which reduces intracellular carbapenem concentrations. While non-CP-CR*Ec* mechanisms are becoming more well understood, there remains limited knowledge of how ESBL-*Ec* strains initially adapt to carbapenem exposures prior to acquiring clinically defined carbapenem resistance.

Perhaps the most successful multi-drug resistant (MDR) *E. coli* lineage of the 21^st^ century is the extraintestinal pathogenic sequence type (ST) known as ST131 [8–11]. Given this lineage’s global dissemination and wide range of observed AMR phenotypes, ST131 has been considered an excellent model for the study of AMR evolutionary trajectories [12–14]. Although there is significant spatiotemporal heterogeneity in the population structure of carbapenem resistant *E. coli* [15, 16], a recent population-based United States surveillance study of carbapenem resistant Enterobacterales found that ST131 was the leading cause of carbapenem-resistant *E. coli* infections with the vast majority having non-CP-CR*Ec* mechanisms [17].

Given the substantial prevalence of ST131 causing non-CP-CR*Ec* infections, we sought to perform a comprehensive assessment of ESBL positive ST131 isolates initial adaptation to carbapenems using a multidisciplinary approach. We determined the initial carbapenem resistance capacity of representative isolates from a large cohort of ST131 bacteremia causing strains as well as comprehensively assessed the long-term evolutionary trajectory of a ST131 strain from the multi-drug resistant associated C2/*H*30Rx subclade under carbapenem selection in distinct experimental evolutionary platforms. Concurrent with data derived from clinical observations of non-CP-CR*Ec* ST131 strains, we found that insertion sequence mediated gene amplification was a significant component to the gradual increase in carbapenem resistance. The convergence of ESBL gene amplification as an initial response within both experimental evolution trajectories in addition to our genomic observations across multiple ST131 subclades suggests that upstream targets activating these cognate responses could be a potential target for reducing progressive resistance to carbapenems and potentially other clinically relevant antibiotic therapies.

## METHODS

### Data collection

ST131 *E. coli* bacteremia events were abstracted from our REDCap database and cross-referenced with MicroBank isolates saved at −80°C in 20% glycerol stocks. WGS sequencing results of *E. coli* bacteremia isolates identified as ST131 from previous studies were included in the study [5, 6, 18–20].

### Antimicrobial susceptibility testing

Clinical isolates were tested for their minimum inhibitory concentrations (MICs) to ertapenem (ETP) and meropenem (MEM) using both gradient strip ETest as well as broth microdilution (BMD) following CLSI M100 (2022) guidelines [21].

### Modified Luria Delbruck assay

A modified fluctuation assay was performed for the purposes of measuring mutation frequency rate in our samples as previously described [22, 23]. Briefly, plate swabs (3-5 colonies) were subcultured in Mueller Hinton Broth (MHB), grown to mid-log phase (∼0.5 OD_600_), and then diluted down to a 0.5 MacFarland standard. Cells were then inoculated at baseline to ∼100 cells/well and grown in MHB for 3 hours at 37°C in 2 96-well plates (192 wells total). After 3 hours elapsed, carbapenem was added at ∼0.75× MIC and samples were grown for 20 hours at 37°C. Ten μL was aliquoted from a random well prior to carbapenem exposure and serially diluted out on MHA plates for colony count enumeration. The BioTek Synergy HTX Multimode plate reader was used to measure optical density (600 nm) with Gen5.3.10 software. Wells with OD_600_ >=0.1 measurements after 20 hours were classified as mutants. Ten μL of culture from positive wells were then re-inoculated with ∼0.75× carbapenem MIC to confirm mutant growth. For samples that had detectable positive growth, up to four mutants were randomly selected for ETP MIC determination using broth microdilution as previously described above and stored at −80C for future genomic characterization as described below.

### Microfluidic system and flask transfer experimental evolutionary platform protocols

The C2*H*30Rx strain MB1860 [5] was passaged using two distinct experimental evolutionary platforms. The novel microfluidic system (MFS) methodology has been published [24]. Briefly, samples are grown overnight in LB, and samples are standardized to 2.5e7 CFU/mL prior encapsulation into microdroplets. Additionally, aqueous (30 μL/min) and oil (100 μL/min) phase flowrates were adjusted to achieve a desired lambda = 10 (cells/microdroplet). After incubation overnight at 37°C, the cells were decapsulated, quantitated and then re-encapsulated at ∼10 cells/microdroplet again to complete one cycle of experimental evolution. ETP concentration was gradually increased to produce a very weak selection gradient from 0 to 0.5 ug/ml ETP over 53 days. For the standard, batch culture, flask transfer protocol (FTP), individual MB1860 colonies that were streaked out on LB agar were grown up in triplicate overnight in LB broth, incubated at 0.5×, 1×, 1.5×, 2× ETP MIC and incubated at 37°C 250 RPM. Next day isolates with highest observable growth were then passaged at until growth was observed at the end point ETP concentration (32 μg/mL ETP). Each MFS passage/iteration was approximately 48 hours (*e.g.,* Day 22 isolates would be equal to the 11^th^ passage/iteration) whereas each FTP passage was equal to 24 hours. Daily populations as well as end-point isolates (EPIs) collected from colonies streaked out from population with last ETP exposure were collected for further characterization described below.

### DNA-Seq

Mutants collected from our modified fluctuation assay of ST131 subclades (n=52), and daily MB1860 population isolates collected from experimental evolution platforms (n=64) respectively were collected for Illumina short-read sequencing. Samples underwent genomic DNA extraction and Illumina short-read, paired-end sequencing DNA library prep as previous described [6]. A total of four samples [MB6054 (n=4); MB10029 (n=1)] from the modified fluctuation assay were censored due to paired-end read contamination leaving us with a total of 48 samples available for mutant analysis.

### RNA-Seq

Daily MB1860 population isolates from both experimental evolution platforms (*i.e.*, FTP and MFS) were selected for RNA-Seq. In particular, we selected daily populations that were exposed to ∼1× ETP MIC = 0.0625 μg/mL (*i.e.*, FTP Population 1, Day 2 isolates and MFS Population 1, Day 26 isolates; we collectively refer to these isolates as ‘experiment’) as well as passaged isolates collected from the same founder population and same day with no ETP exposure from each respective population (i.e., FTP-P1-D2 and MFS-P1-D26 without ETP exposure). Three technical replicates from each group were grown to mid-log phase (OD600 = 0.5 ± 0.05) with the experimental isolates grown in LB broth supplemented with 0.03 μg/mL ETP whereas control isolates were grown without ETP exposure. Three mL of each sample was treated with RNAprotect (QIAGEN) following manufacturer’s instructions, and pellets were stored at 80°C until each sample and respective replicate was ready for RNA extraction. RNA was extracted using the QIAGEN RNeasy Mini Kit using manufacturer’s instructions. RNA-seq was performed on an Illumina NovaSeq6000 instrument.

### Western blot analysis

Immunoblot analysis of outer membrane porins has been described previously [25]. Briefly, samples are grown to mid-log phase and then standardized by OD600. Samples are then boiled and subjected to electrophoresis on SDS polyacrylamide gel supplemented with 4M urea. Anti-rabbit antibodies targeting OmpC/OmpF/OmpA have been previously validated and reported [26, 27].

### Comparative Genomics

A total of 183 ST131 *E. coli* paired-end short-reads that were sequenced from previous molecular epidemiology projects [5, 6, 18–20] were aligned to MB1860 (RefSeq #: NZ_CP049085.2) using the Snippy-v4.6.0 variant call pipeline (Seemann, T. snippy GitHub: https://github.com/tseemann/snippy). Core genome alignment file was used to infer a recombination-free maximum likelihood phylogenetic tree using UFBoot2 (1000X) and ModelFinder with IQTree-2-v2.0.3 [28] and Gubbins-v3.2.1 [29]. The filtered polymorphic sites output from Gubbins was used as input to the hierBAPS (rhierbaps-v1.1.4) function to reveal subclade structure [30] using default parameters and level 2 as subclade assignment. Comprehensive ST131 subclonal genomic characterization was performed using the ST131typer command-line tool [31]. AMRFinderPlus (v3.11.26) was performed to determine both AMR and virulence factors identified in both short- and long-read assemblies using database version 2024-01-31.1. Convict-v1.0 was used to estimate AMR gene copy numbers as previously described [6].

### Differential Expression Analysis

MB1860 passaged isolates were selected in the MFS and FTP system with and without ertapenem (ETP) selective pressure (passage control - 0 ug/mL; experiment - 0.0625 ug/mL ETP). BBDuk was used to trim Nextera Illumina adapters from RNA paired-end reads using default parameters with modifications [32]. Fast-QC-v0.12.1 [33] was used to perform quality control on trimmed RNA reads. One sample (MFS-P1-E3 = Experimental sample 3) was censored due to low read quality metrics based on Fast-QC output. Trimmed RNA-reads were aligned to MB1860 (RefSeq #: NZ_CP049085.2) using the STAR aligner [34]. Htseq was used for enumerating read counts of uniquely mapped reads [35]. The R package DESeq2 (v1.42.0) was used for differential gene expression analysis [36]. Median of ratios was used to normalize transcripts and log2 fold change were shrunk to stabilize for variance and low read counts. Gene set enrichment analysis was performed on DESeq2 output with the R package ClusterProfiler (v4.10.0) [37].

### Statistics

All statistical analysis was performed in R-v4.3.2 as previously described [6, 19, 20].

### Data Availability

All R Scripts used for this analysis can be found in the Figshare published repository, DOI: 10.6084/m9.figshare.26400901. Whole genome sequencing data has been made publicly available through the NIH BioProject repository PRJNA1141466 and GEO Series accession number GSE273456.

## RESULTS

### The majority of ST131 non-CP-CR*Ec* isolates are detected within the C2/*H*30Rx subclade

There was a total of 184 ST131 bacteremia isolates from 171 unique patients (*i.e.*, 13 cases of recurrence, Table S1). **Figure 1** illustrates the core ST131 genome population structure with the previously designated three major subclones labelled. The vast majority of isolates (87%, 160/184) were from the worldwide disseminated ‘C clade’, often designated as *H*30R [31] due to its strong association with the *fimH30* allele and fluoroquinolone resistance [38]. There were 11 clusters of ST131 clades identified using a hierarchical Bayesian analysis of the core genome as represented by branch tip color (**Figure 1**). Carbapenem resistance was present in 6% (11/171) of unique patient ST131 strains with three patients infected by strains that had phenotypic cephalosporin resistant to non-CP-CR*Ec* conversion (Table S1). Nearly all carbapenem resistant isolates (91%; 10/11) were non-CP-CR*Ec*. Half of all non-CP-CR*Ec* isolates were detected within the cluster 7 C2/*H*30Rx subclade (n=5) with 80% co-harboring *bla*_CTX-M-15_ and *bla*_OXA-1_ as detailed in **Figure 1**. The second most commonly detected ESBL encoding gene detected within non-CP-CR*Ec* isolates was *bla*_CTX-M-27_ (n=3). When characterizing variants across the ten non-CP-CR*Ec* isolates (Table S2), the only genes with disruptive mutations detected across two or more isolates were the outer membrane porin encoding genes *ompC* and *ompF* as well as the transcriptional regulator encoding gene *soxR* which can negatively regulate efflux activity [39, 40].

**Figure 1.**
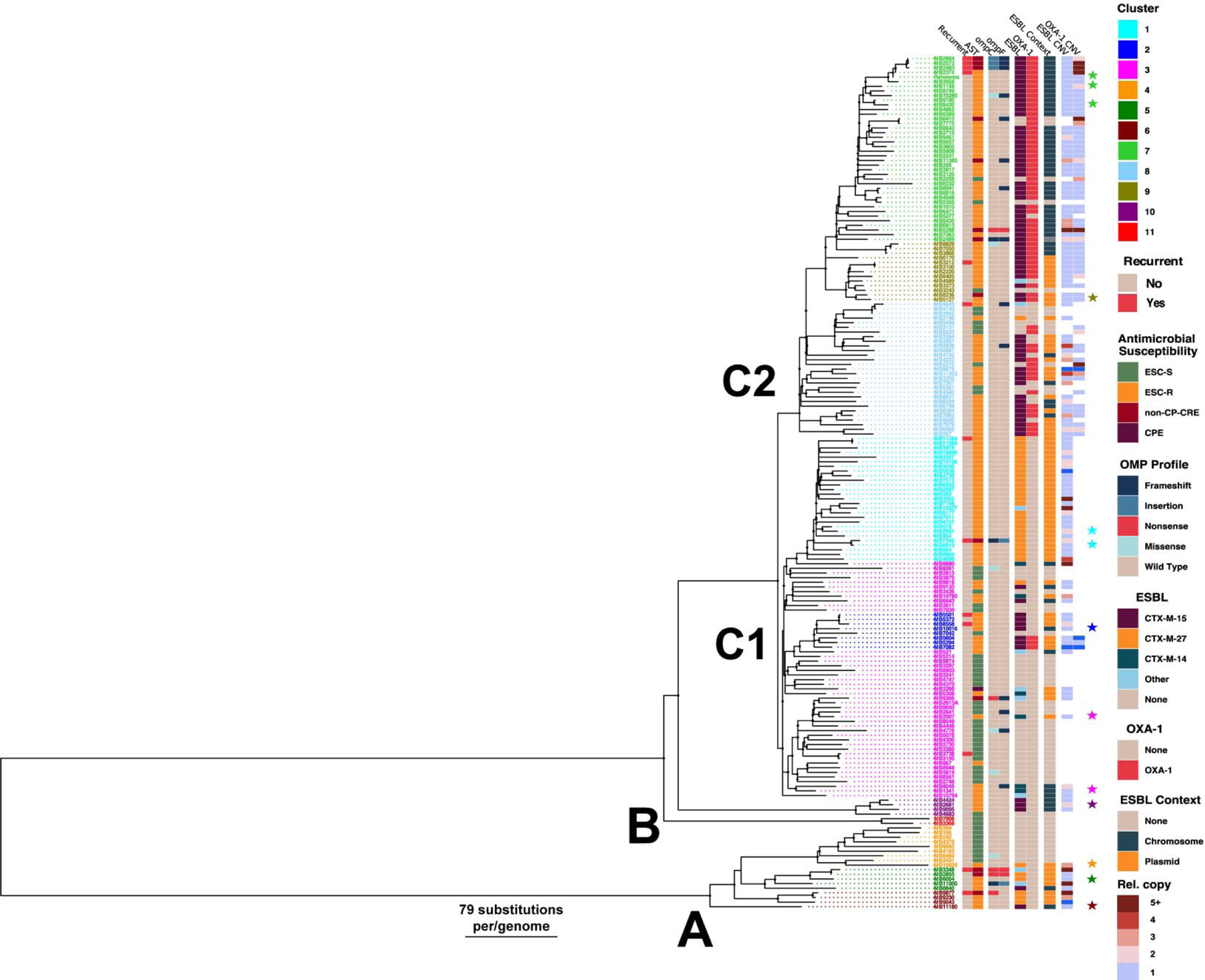
Population structure of ST131 *Escherichia coli* bacteremia isolates. Mid-point rooted, recombination-masked, core-genome inferred maximum-likelihood phylogeny of ST131 *E. coli* (n=184) collected from 2013 to 2020 at our institution. The three major subclones with clade C split into C1 and C2 are labelled on the dendrogram. Terminal branch tips are colored by sub-cladal structure (n=11) as identified using a Bayesian analysis of population structure. Columnar metadata are labelled with corresponding categories presented in figure legend on the right. Thirteen ST131 isolates from nine sub-clades that were selected for carbapenem mutation frequency analysis are labelled by stars that appear to right of columnar metadata.

### ESBL positive ST131 bacteremia isolates increase ESBL gene copy number in response to initial ertapenem exposure across multiple ST131 subclade backgrounds

To gain insight into the initial ESBL-*Ec* adaptation to carbapenems, we selected thirteen carbapenem-susceptible ESBL-positive ST131 isolates from differing subclade backgrounds and measured mutation frequency following exposure to ertapenem (ETP) using a fluctuation assay. We observed no mutants above the limit of detection for three ESBL-negative ST131 isolates (Table S3). All thirteen ESBL positive isolates had detectable mutation frequencies ranging from 106 to 12854 mutants per 10^8^ cells (**Table 1**). A strong positive linear relationship was detected between initial ESBL gene copy number in the parental isolate and subsequent mutation frequency calculated (*r* = 0.88; *P*-value < 0.001). We selected a subset of ST131 isolates (n=6) to measure their meropenem (MEM) mutation frequency and detected a significantly narrower range from 69 to 381 mutants per 10^8^ cells suggesting a smaller adaptive mutation supply for MEM in contrast to ETP (Table S4).

**Table 1.** Carbapenem Fluctuation Assay Results for Select ST131 ESBL Positive Strains.

|  |  |  |  |  | Ertapenem (ETP) Fluctuation Assay Results |  |  |  |
| --- | --- | --- | --- | --- | --- | --- | --- | --- |
| Sample | Clade<br>(sub-level) | ESBL | ESBL<br>Context | P<sub>0</sub> ESBL<br>CNV | P<sub>0</sub> ETP<br>MIC<br>(μg/mL)<sup>a</sup> | P<sub>0</sub> Mut Freq<br>Mut/10<sup>8</sup><br>Cells<sup>b</sup> | P<sub>1</sub> ETP<br>MIC<br>(μg/mL)<sup>a</sup> | P<sub>1</sub> ETP<br>Fold<br>Change (×) |
| MB10029 | A (4) | CTX-M-27 | plasmid | 2.97 | 0.094 | 12854 | 0.25 | 3 |
| MB6054 | A (5) | CTX-M-27 | Plasmid | 1.42 | 0.063 | 564 | 0.25 | 4 |
| MB11180 | A (6) | CTX-M-15 | chromosome | 0.93 | 0.5 | 967 | 1.5 | 3 |
| MB2681 | B0 (10) | CTX-M-15 | chromosome | 1.69 | 0.25 | 2879 | 0.5 | 2 |
| MB10016 | C1 (2) | CTX-M-15 | chromosome | 1.01 | 0.094 | 106 | 0.5 | 5 |
| MB1341 | C1 (3) | CTX-M-14 | chromosome | 1.13 | 0.25 | 998 | 0.5 | 2 |
| MB2007 | C1 (3) | CTX-M-14 | plasmid | 1.23 | 0.063 | 117 | 0.5 | 8 |
| MB2951 | C1-M27 (1) | CTX-M-27 | plasmid | 1.67 | 0.094 | 6874 | 1.0 | 11 |
| MB6910 | C1-M27 (1) | CTX-M-27 | plasmid | 1.47 | 0.063 | 749 | 0.25 | 4 |
| MB1159 | C2 (7) | CTX-M-15 | chromosome | 1.06 | 0.13 | 1368 | 1 | 8 |
| MB1860 | C2 (7) | CTX-M-15 | chromosome | 0.93 | 0.25 | 2242 | 1 | 4 |
| MB8420 | C2 (7) | CTX-M-15 | chromosome | 0.85 | 0.25 | 2031 | 1 | 4 |
| MB5127 | C2 (9) | CTX-M-15 | plasmid | 1.37 | 0.5 | 1149 | 1 | 2 |

$P_0$ = ancestral clinical strain; $P_1$ = mutant progeny strain
<sup>a</sup>MICs are reported as the median of two or more repeat experiments performed in duplicate
<sup>b</sup>Mut Freq = Mutation Frequencies reported as the mean number of mutants per 10<sup>8</sup> cells (n = 3 replicates)

We characterized three to four mutants per parental strain and found that the median ETP MIC fold change was elevated and generally similar across subclades with the exception of a statistically significant difference between B0 and C1-M27 subclades (**Fig 2A**). When focusing on mutant ESBL gene fold changes relevant to parental strain, there was a range across ST131 subclades with the C2/*H*30Rx mutants having the greatest increase (median = 3.7×; interquartile range (IQR) = 1.1×) and A subclade mutants having only a minimal increase (median = 1.1×; IQR = 1.1×) (**Fig 2B**). Variant calling analysis of 48 mutant strains indicated that the majority did not have SNPs detected relative to parental isolates (median = 0; interquartile range (IQR) = 1) (Fig S1). Twenty-three percent (11/48) had a variant that would be predicted to affect carbapenem susceptibility (Table S5) with the majority (n=8) having a predicted effect on outer membrane porin integrity including *ompC*, *ompF*, and *surA,* the latter which encodes a chaperone protein that is responsible for proper folding of outer membrane proteins [41, 42]. When comparing ESBL gene amplification stratified by variant status, mutants that had a variant of interest (*i.e.*, mutation predicted to affect ertapenem MIC) had statistically lower ESBL gene fold changes (median = 0.9×; IQR = 0.2×) compared to isolates without a variant detected (median = 3.3×; IQR = 2.7×) (Wilcoxon Rank-sum Test *P*-value = 0.0001) (**Fig 2C-D**). Variants of interest were mostly detected in A, C1, and C1-M27 subclades (**Fig 2B**) in contrast to C2 subclade where ESBL gene amplification appeared to be the predominant genomic adaptation to ertapenem exposure.

**Figure 2.**
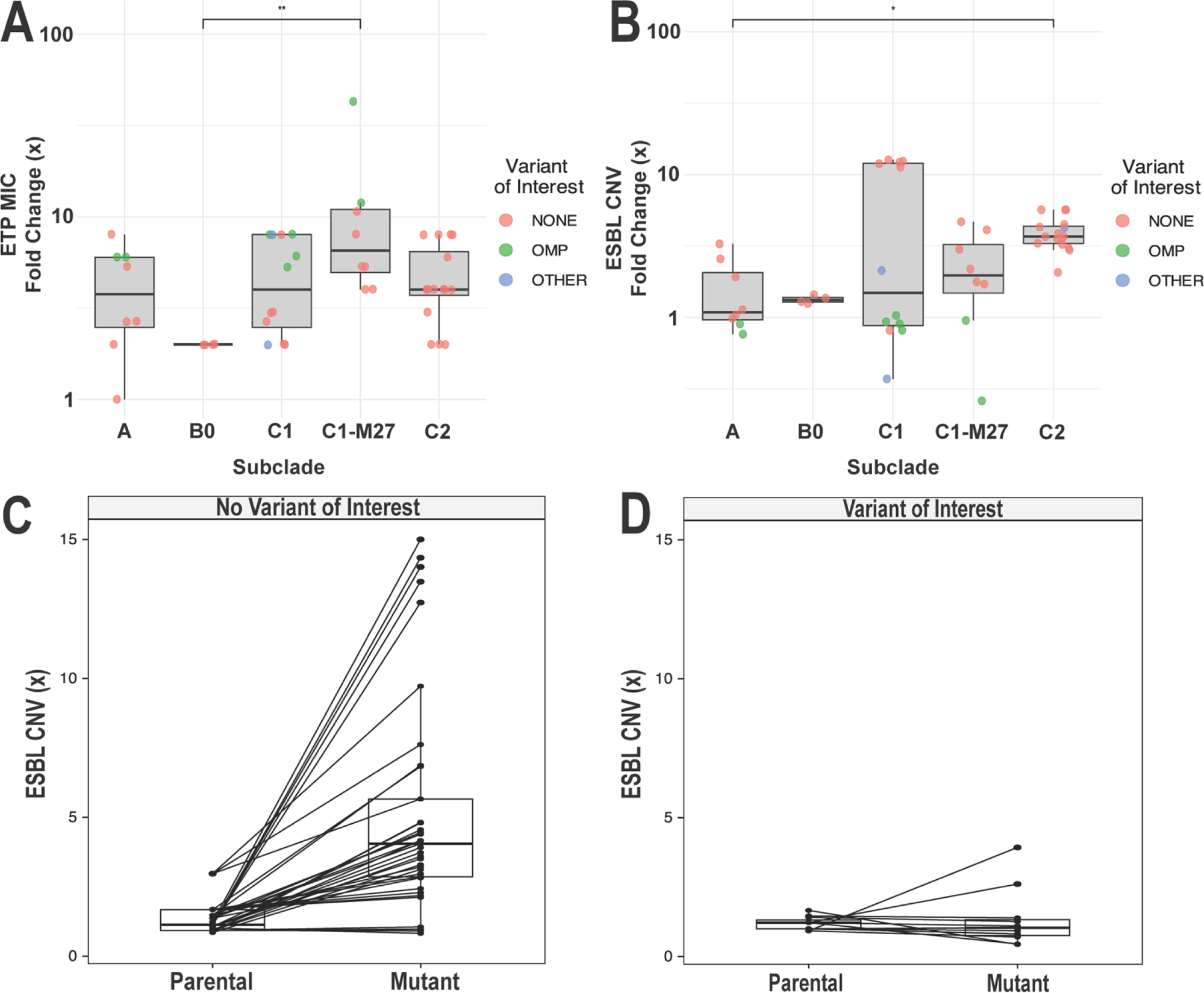
ST131 Fluctuation Assay Mutant Genomic Characterization. **(A)** ETP MIC fold change (×) and **(B)** ESBL gene fold change (×) across five predominant subclades of ST131. Note that the y-axis in each respective subpanel is presented in log10 scale without data transformation. Variant status is labelled by color as presented in figure legend. ESBL copy number variant (CNV) between parental (*i.e.*, carbapenem naïve) and progeny mutant (*i.e.*, positive growth following ETP exposure) strain stratified by isolates with no variant of interest **(C)** versus strains with a variant of interest detected **(D)**. Dunn multiple-comparison ** = *P*-value < 0.01; * = *P-*value < 0.05

### The C2/*H*30Rx strain MB1860 initial response to ertapenem exposure is amplification of IS*26*-mediated pseudocompound transposons (PCTns)

We passaged MB1860 in the presence of ETP within both standard, serial flask transfer protocol (FTP) and microfluidic system (MFS) platforms as a means to model ‘fast’ vs. ‘slow’ adaptation to ertapenem exposure respectively [24]. We studied the MB1860 strain since it belonged to the subclade with the largest number of non-CP-CR*Ec* isolates (*i.e.*, C2/*H*30Rx) and because it contains a chromosomally located pseudocompound transposon (PCTn) harboring *bla*_CXT-M-15_ and *bla*_OXA-1_ genes, which we formerly described as Tn*MB1860* [5]. All three biological replicates from the FTP experiments developed high level (*i.e.,* ETP MICs ≥ 32 μg/mL) ertapenem resistance (ETP-R) within 9 – 12 days (**Fig. 3A**). Conversely, it took 53 days to adapt MFS populations to ETP concentrations equal to 0.5 μg/mL (**Fig. 3B**), which may reflect the stronger selection and slower adaptation within MFS environment.

**Figure 3.**
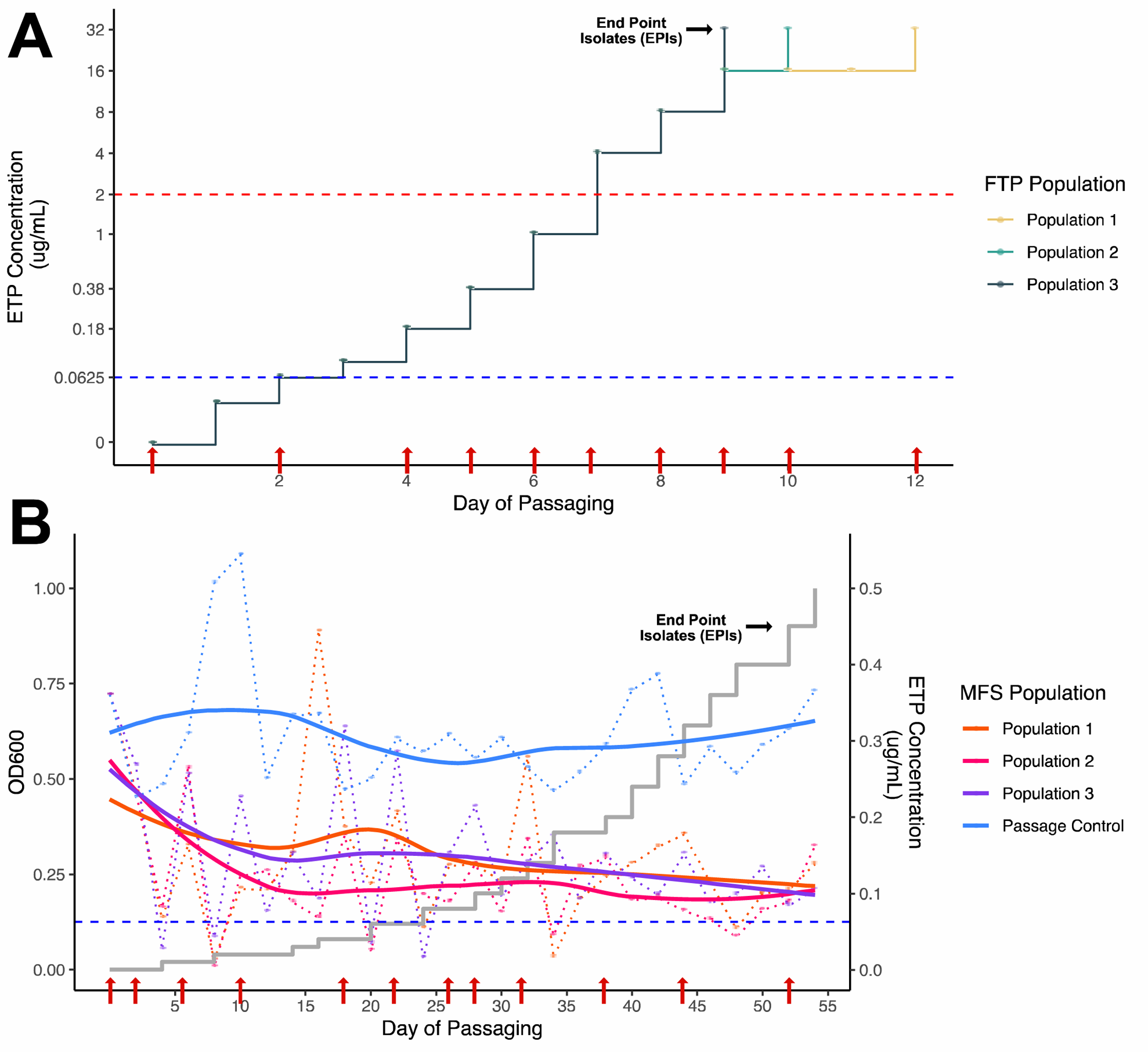
Growth of MB1860 with increasing ETP concentration gradients. **(A)** FTP populations (n=3) exposed to stepwise gradient increase in ETP concentrations with indication of positive growth (OD600 > 0.5) at each respective day. Note that Population 1 – 3 stepwise growth followed comparable trajectories up to day 9 of passaging **(B)** MFS populations (n=3) with OD600 measurements indicated on the left y-axis and ETP concentration stepwise increments (grey solid line) indicated on right y-axis. Dotted lines indicate variance in OD600 measurements over time with each population color-coded in legend. A fitted loess curve (solid color-coded lines) for OD600 measurements of MFS populations is juxtaposed across the line graph. Blue horizontal dotted line indicates ancestral MB1860 ETP MIC (0.0625 μg/mL). Red horizontal dotted line **(A)** indicates CLSI carbapenem resistance breakpoint [21]. Red arrows indicate WGS sampling timepoints.

Consistent with the data from the mutation frequency analysis, for both the FTP and MFS platforms, we detected *bla*_OXA-1_ and *bla*_CTX-M-15_ amplifications at ETP concentrations as low as 0.06 and 0.01 μg/mL in the FTP and MFS systems respectively (**Fig. 4AB**). Genome-wide analysis of daily population isolates exposed to 1× ETP MIC (FTP Day 2, MFS Day 23) showed that, for both FTP and MFS platforms, gene amplification was focused at PCTn regions (Fig. S2A), with a high degree of modularity with respect to IS*26* elements being amplified (Fig. S2B). For example, in FTP population 1 and population 3 (we refer to as ‘FTP-P1’ and ‘FTP-P3’ respectively), we consistently observed amplification of a nearly 60 Kb region (Fig. S2B) starting with the 5’ insertion of the MB1860 PCTn into the *cirA* gene and downstream of another IS*26* element that truncated a DUF encoding gene (**Fig. 4A**). In contrast, for the MFS populations, gene amplification occurred only in the ∼13 Kb MB1860 PCTn region for MFS-P1 isolates, the whole 60 Kb region for MFS-P2 isolates, and initially the 60 Kb until the final MFS-P3 day 52 (*i.e.*, MFS-P3-D52) isolate had PCTn amplification only (Fig. S2B and **Fig. 4B**). For all isolates, we observed significant amplification of *bla*_CTX-M-15_, up to 20x, whereas *bla*_OXA-1_ was inconsistently amplified both over time, amongst isolates, and between platforms (**Fig. 4**). Differential amplification patterns could be observed across each of the experimental platforms as well as each of the independent populations, highlighting the dynamic nature of PCTns mediated by IS26 elements (Fig. S3A-E).

**Figure 4.**
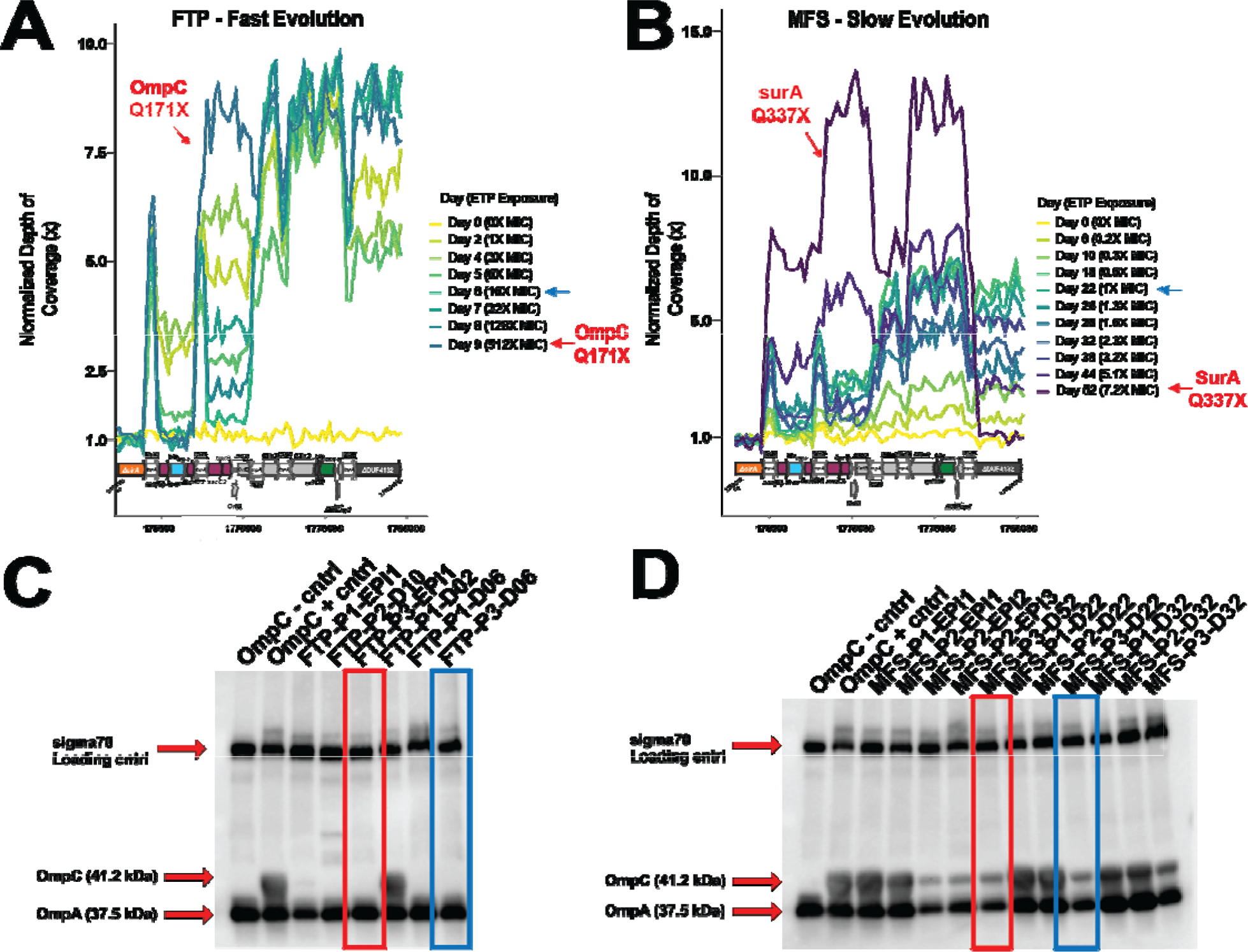
Amplification of the MB1860 pseudocompound transposon and Western Blot analysis of outer membrane porins (OMP) in FTP and MFS experiments. **(A)** FTP and **(B)** MFS normalized coverage depth of MB1860 PCTns indicating amplification of beta-lactamases *bla*_OXA-1_ (blue open reading frame) and *bla*_CTX-M-15_ (green open reading frame) as presented on X-axis. Red arrows indicate where nonsense mutations were detected corresponding with each particular day. FTP Population 3 (FTP-P3) and MFS-P3 isolates are illustrated here as representatives of their respective evolutionary trajectories. **(C)** FTP and **(D)** MFS immunoblot analysis of outer membrane porin C (OmpC) across select daily population and end point isolates (EPIs). ‘OmpC – control’ is previously characterized OmpC negative strain MB9877 [6] and ‘OmpC + control’ is the positive control, parental MB1860 strain. Red and blue boxes correspond to red and blue arrows respectively in **(A)** and **(B)**, highlighting strains of interest.

Independent of experimental evolutionary platform, there were only a small number of genetic variations, the majority of which appeared after prolonged, increasing ETP exposure (Fig. S3A-F and Table S6). For the FTP strains, variants of interest included nonsynonymous mutations in *ompC* (outer membrane porin C), *ompF* (outer membrane porin F), and *purM*, which encodes a purine metabolism gene (Fig. S3D-E). Other mutated genes with common functions including *efp* and *epmA*, which encodes proteins involved in translation elongation, as well as *wecA* which encodes a protein important to outer membrane composition. For the MFS platform, we identified no variants of interest in daily populations with the exception of a nonsense mutation in *surA* from P3 isolates and end point isolates (EPIs) collected from the last founder population exposed to ETP (Fig. S3F); nonsynonymous mutations were present in genes encoding proteins involved in outer membrane composition such as the histidine kinase regulator of OmpC/F expression, EnvZ (Pop 1), and the periplasmic chaperone protein responsible for efficient assembly of OmpC/F, SurA (Pop 2 and 3). Similar *surA* mutations, which would be predicted to disrupt OmpC/F production, were identified in a subset of our mutant fluctuation assay results described above during initial ETP adaptation.

In light of the recurrent finding of outer membrane protein variation in all our assays, we tested for the presence of the key outer membrane protein OmpC using Western immunoblot as shown in **Fig. 4CD**. For the FTP passaged strains, we only detected OmpC at day 2 whereas no OmpC was detected by day 6 even in the absence of genetic mutations affecting *ompC* detected from this daily population (**Fig. 4C**; blue box). No OmpC was identified in any of the FTP EPIs consistent with the *ompC* mutations detected in these strains exposed to high ETP concentrations (**Fig. 4C**). For the MFS isolates, we consistently detected OmpC production; however, the OmpC production was decreased in the MFS-P3-D52 isolate (**Fig. 4D**; red box) corresponding to when a *surA* nonsense mutation was detected as well as MFS-P3-D22 isolate prior to any detected genomic changes (**Fig. 4D**; blue box). This is consistent with reduced expression of OmpC being an important adaptation in response to higher concentrations of ETP.

### Identification of consistent impact of IS26 on the MB1860 transcriptome in response to ETP exposure

To identify genes with altered transcript levels following ETP exposures in both experimental evolutionary systems, we performed RNA-seq on cells grown with and without ETP exposure (**Fig. 5**). A Pearson correlation coefficient heatmap of differential expression data is presented on Fig. S4. Of the 24 genes with significantly differentially expressed transcript levels following ETP exposure in both platforms (defined as at least absolute value log2 fold change > 1 and adjusted *P*-value < 0.05), 12 were genes that were harbored within the MB1860 PCTn including *bla*_OXA-1_ (**Fig. 5A**) and *bla*_CTX-M-15_ (**Fig. 5A-B**) genes (Table S7). Indeed, of all the genes with increased transcript levels in both systems, *bla*_CTX-M-15_ had the highest (in the FTP) and 2^nd^ highest (in the MFS) normalized read counts indicating the marked transcriptional output dedicated to *bla*_CTX-M-15_ following ETP exposure (Table S7). When performing gene set enrichment analysis, the one activated gene set activity conserved across both platforms was ‘transposase activity’ (**Fig. 5C-D**). The gene set fraction was <30% for each group and largely confined to ISs (e.g., IS*26*) that were bracketed by the MB1860 PCTn (Table S7).

**Figure 5.**
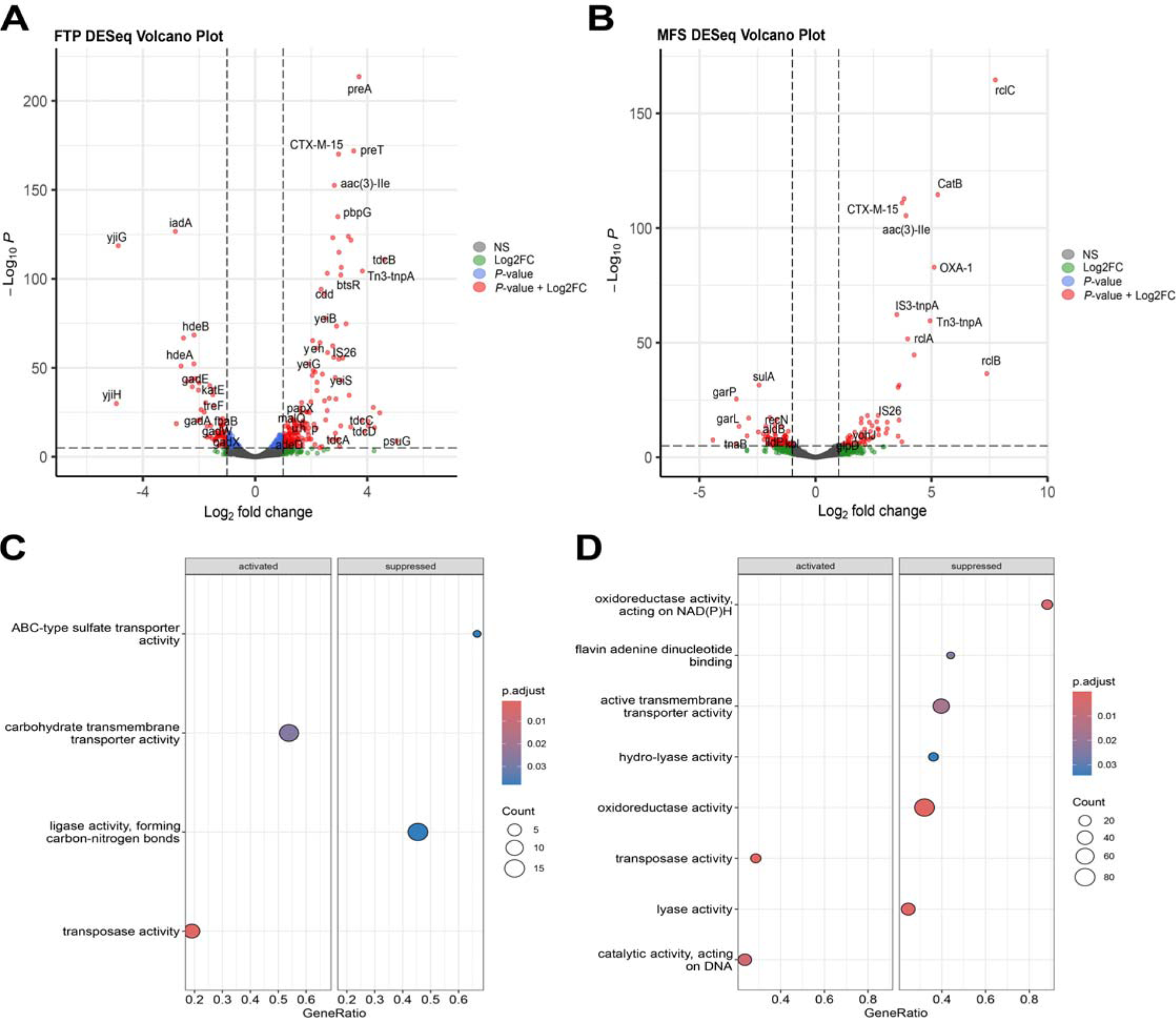
Transcriptome analysis of MB1860 daily population isolates in FTP and MFS platforms. Differential expression analysis was conducted on daily population isolates (FTP Population 1, Day 2 isolates and MFS Population 1, Day 26 isolates) and exposed to 0.5× ETP MIC (*i.e.*, 0.03 ug/mL ETP) compared to passage controls without ETP exposure. Volcano plots of differentially expressed genes in **(A)** FTP and **(B)** MFS platforms respectively with genes of interest identified. Dotted lines demarcate genes that have an adjusted P value cut-off of 1e-6 and Log2 fold change = 2 with red dots indicating statistically significant results with absolute log2 FC values >2. Gene set enrichment analysis of **(C)** FTP and **(D)** MFS platforms respectively with activated v. suppressed gene sets. Gene ratio indicates total fraction of genes belonging to that particular gene set with statistically upregulated or downregulated expression and count labelled in legend respectively.

## DISCUSSION

There has been recent research characterizing the predominant carbapenem resistance mechanisms by which *E. coli* either acquire carbapenemases through horizontal gene transfer or increase production of β-lactamase encoding genes and concurrently reduce cellular permeability [43]; however, there has remained a gap in knowledge with regards to how high-risk ESBL-*Ec* adapt to initial carbapenem exposure. Our study objective was to comprehensively investigate the genomic and transcriptomic responses to initial carbapenem exposure using ETP selection with the successful ExPEC ST131 lineage as a model to determine evolutionary pathways. Using multiple complementary strategies, we found that IS-mediated gene amplification is the predominant ST131 adaptation to ETP exposure, which can potentially progress to non-CP-CR*Ec* development.

A key finding of our study was the identification of IS-mediated amplification of ESBL encoding genes as a primary ESBL ST131 adaptive response, particularly for C2/*H*30Rx strains (**Fig. 2**). Our comprehensive multiomic analysis permitted us to look at multiple mechanisms of carbapenem resistance such as increased efflux activity and transpeptidase modifications that may affect carbapenem binding activity. Nonetheless, our comparative genomics analysis of mutants recovered from our fluctuation assay indicated ST131 mutants consistently either initially responded to ETP exposure by increasing ESBL gene copies or disrupting porin production (**Fig. 2CD**). Multiple studies have shown how *E. coli* and other *Enterobacterales* spp. modulate their outer membrane permeability to decrease carbapenem influx into the periplasm and concurrently increase expression of ESBL or pAmpC encoding genes [7, 44–47]. Nevertheless, both reduced porin-mediated permeability [48] and gene amplification in the absence of selection [49, 50] have been shown to impart significant fitness costs potentially limiting transmission. A recent United Kingdom comparative genomics study has demonstrated in a case series of a clinical *E. coli* isolate that despite high PCTn-mediated chromosomal amplification of *bla*_TEM_, this particular isolate did not incur significant fitness costs [51]. Furthermore, recent studies from different geographic locales [52–54], have found consistent with our study, a high prevalence of the ST131 C2/*H*30Rx subclade with stable chromosomal PCTn incorporation harboring *bla*_CTX-M-15_ gene in the absence of IncF type plasmid carriage. These results suggest chromosomal integration may impart a survival benefit to ameliorate gene amplification incurred fitness costs that arise from plasmid-mediated *bla*_CTX-M-15_ amplifications. However, increased plasmid-mediated amplification of *bla*_CTX-M-27_ with elevated ETP MIC fold changes in C1-M27 mutant progeny (**Fig. 2AB**) illustrates the potential for multiple, successful evolutionary pathways towards ETP-R development.

That porin mutation imparts a significant fitness cost was reflected in the complex pattern by which the C2/*H*30Rx MB1860 strain altered porin function as detected in our experimental evolution platforms. In both the FTP and MFS platforms, decreased OmpC production at the protein level was not detected until several days after ESBL gene amplification at a time when no genomic mutations were detected (**Fig. 4**). The amount of outer membrane porin protein content in the outer membrane can be affected at the gene (*e.g.,* insertions/deletions [6]), transcription (e.g. two-component signal transduction systems, EnvZ/OmpR [7, 55]), post-transcriptional (e.g. MicF [56]), and even post-translational level (e.g. chaperone proteins like SurA [41]). We did not identify any significant difference in porin gene transcript levels following ETP exposure, but this analysis was only performed at a single time point and thus cannot entirely exclude a transcriptional level explanation for the initial OmpC protein changes; furthermore, differential expression analysis cannot measure changes in sRNA transcripts. For the MFS platform, we did not identify *ompC* or *ompF* mutations, but rather there were mutations in *envZ* and *surA* which was reflected in the decreased but still detectable amount of OmpC protein. This suggests that the fitness cost imparted by *ompC* or *ompF* mutation was too severe under the MFS conditions to become stable in the population suggesting weaker alleles were more successful at low ETP concentration *e.g.,* weaker alleles were sufficient to balance fitness costs against benefits at lower concentrations of ETP. Conversely, in the more nutrient rich FTP platform, inactivating mutations in *ompC* and *ompF* became detectable once ETP exposed passaged populations of MB1860 crossed the ETP-R breakpoint, consistent with these two necessary mechanisms contributing to non-CP-CR*Ec* development observed in our clinical isolates. Whereas frameshifts in *ompC* and/or *ompF* genes are typically stable, it is likely that the initial decrease in OmpC protein levels observed as MB1860 moved towards the carbapenem CLSI breakpoints in the FTP platform was achieved via a reversible process, such as transcriptional or post-translational changes. This reversibility in turn could help explain the recently described category of “Unconfirmed CRE” in which strains which initially tested as carbapenem resistant at a local laboratory did not have the same phenotype upon repeat analysis [17].

Our study has a few limitations including small sample size and a limited characterization of other evolutionary trajectories under alternative carbapenem drug exposures. Nevertheless, our results indicate a robust response to ertapenem selective pressure across all ESBL positive ST131 subclades we tested with repeat measures of initial genomic (*e.g.*, ESBL amplification or Omp mutation) and transcriptomic (*e.g.*, transposase activity) responses, which were consistent with mechanisms we detect in MEM-R clinical isolates [6]. Future work will be necessary to determine upstream targets of gene amplification and how initial carbapenem adaptations contribute to non-canonical resistance mechanisms such as heteroresistance in the clinical setting [57–59].

In summary, we have used experimental evolutionary platforms with complementary multiomic approaches to comprehensively assess the initial adaptation and progression of carbapenem resistance within ST131 *E. coli* bacteremia isolates. Our findings support unique evolutionary trajectories within ST131 subclades with the predominant ST131 C2/*H*30Rx subclade initially responding to carbapenem selection through IS*26*-mediated amplification of ESBL encoding genes in the absence of detected genomic changes affecting outer membrane permeability. Given the emerging understanding of the role of gene amplification in resistance development to multiple classes of antibiotics, increased understanding of gene amplification regulation could lead to new preventative or treatment strategies.

## Acknowledgments

We would like to thank the MDACC clinical microbiology lab for all their hard work in identifying, handling, and transferring these pathogenic strains of interest to us for our research projects. The authors acknowledge the support of the High-Performance Computing for research facility at the University of Texas MD Anderson Cancer Center for providing computational resources that have contributed to the research results reported in this paper.

## Ethical statement

This study has received ethical approval through the University of Texas MD Anderson Cancer Center (MDACC) Institutional Review Board (Protocol ID: PA13-0334).

