## Supplemental Figures for "Comprehensive Assessment of Initial Adaptation of ESBL Positive ST131 *Escherichia coli* to Carbapenem Exposure"

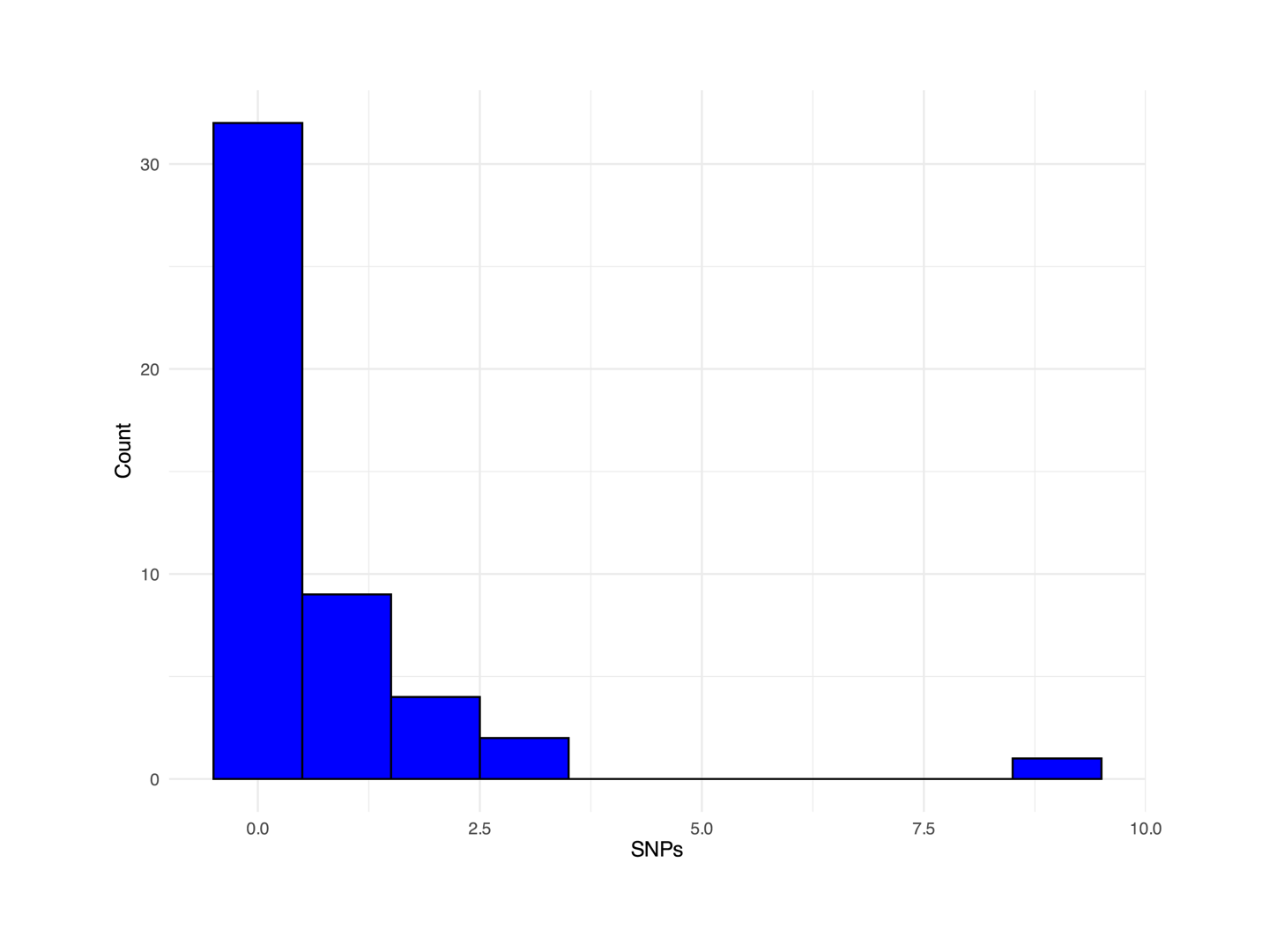

**Fig. S1. Histogram of SNPs detected across 48 fluctuation assay mutants.**

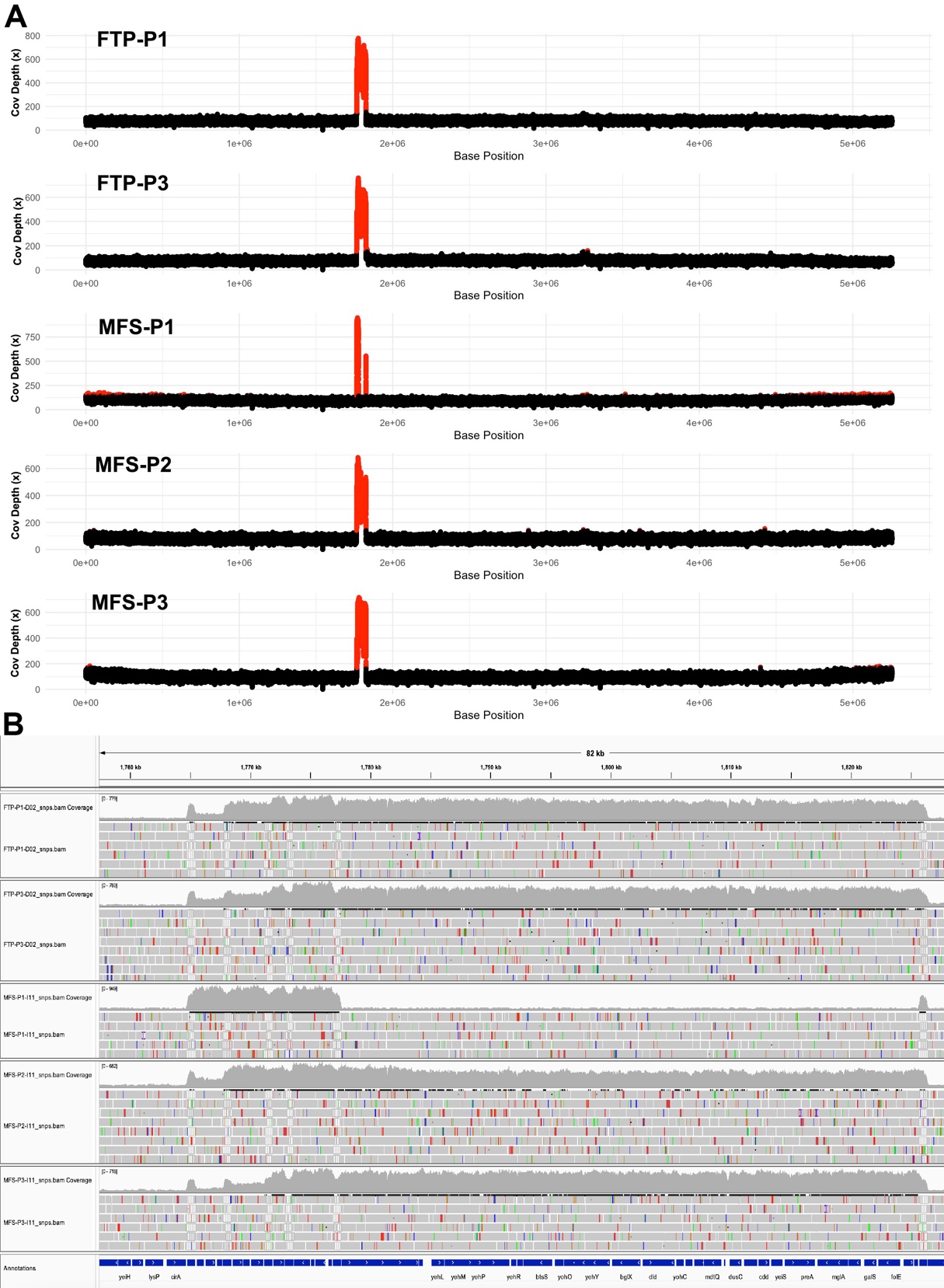

**Fig S2. Chromosome-wide coverage depth for FTP and MFS daily populations. (A)** Genome wide coverage of MB1860 (5 Mb) of FTP and MFS daily populations collected at 1x ETP MIC exposure (Day 2 and Day 22 respectively). Red regions indicate where coverage depth exceeded mean coverage + 1.5 standard deviations. Large red peak for each population corresponds to MB1860 PCTn **(B)** Coverage focused on region corresponding to MB1860 PCTn that is notably increased in **(A)**.

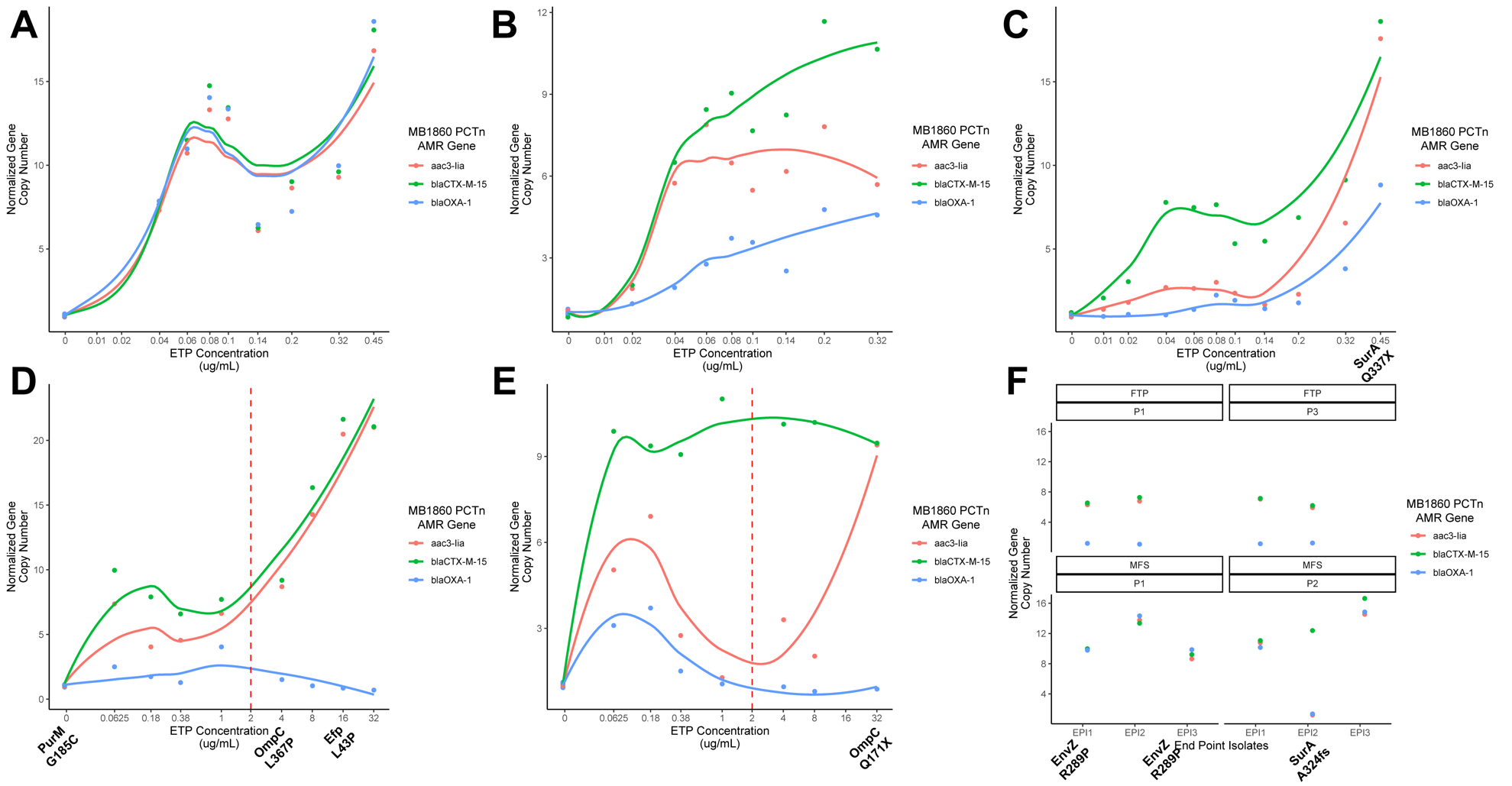

**Fig. S3. Copy number variation of AMR genes located within MB1860 pseudocompound transposon from daily population and end-point isolates. (A-C)** MFS populations (P1-3 respectively) and **(D-E)** FTP populations (P1 and P3 respectively) with variants of interest indicated at ETP exposure they were first detected. Loess curves are included to indicate trend of AMR encoding gene copy number trends. Vertical dotted red line in **(D)** and **(F)** indicate CLSI breakpoint for ETP-R. **(F)** Copy number variation of end point isolates (EPIs) selected from the daily population with last ETP exposure. Note that FTP P3 daily population and MFS P3 EPIs were not available for sequencing respectively.

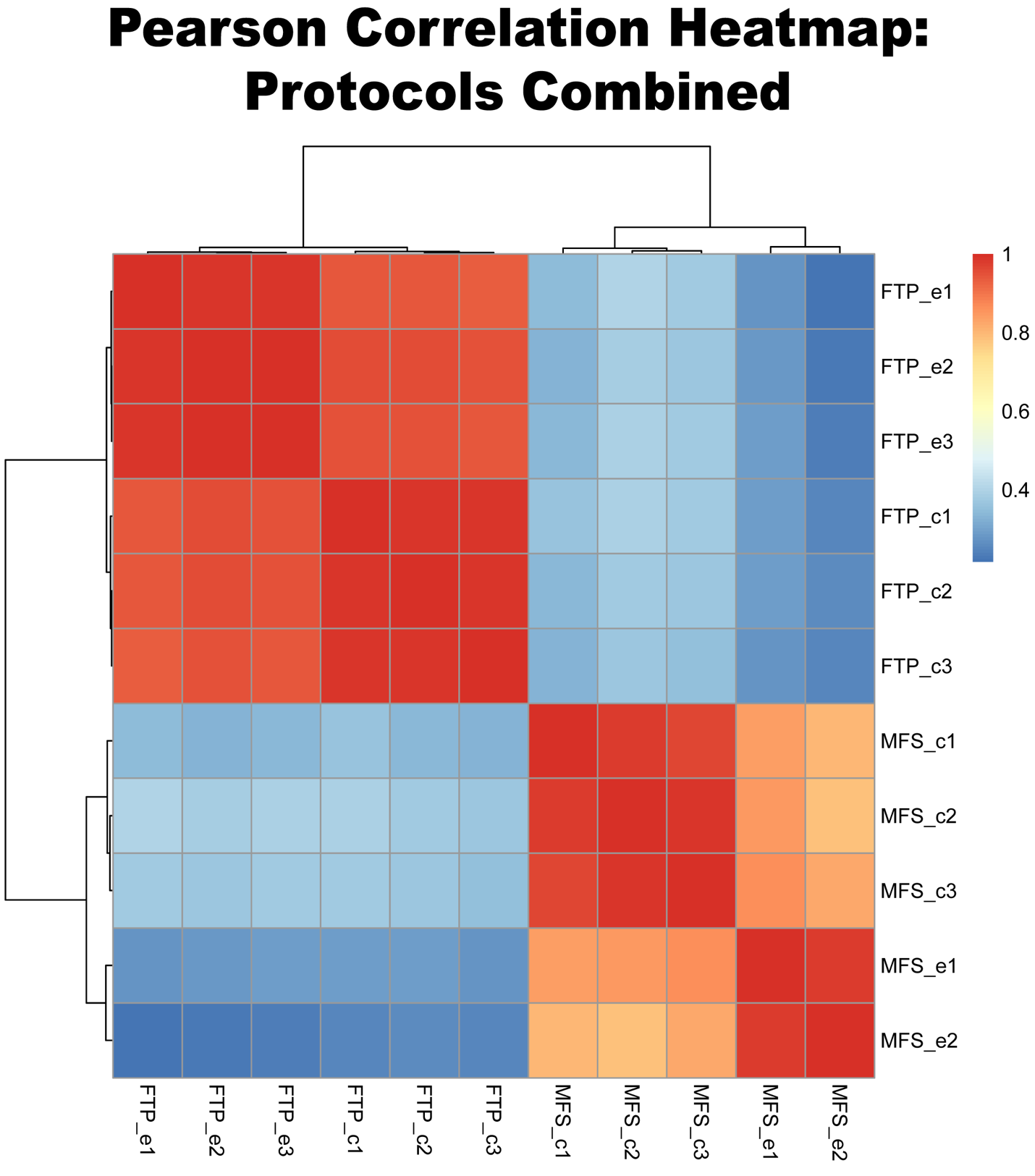

**Fig S4. Correlation heatmap of differential expression observed in FTP and MFS platforms.** Suffix following FTP or MFS with ‘e’ or ‘c’ indicates ‘experiment’ or ‘control’ respectively with replicate number.
